# Chromosome-scale assembly of the *Cupressus sempervirens* genome unravels new insights into the evolutionary history of conifers

**DOI:** 10.64898/2026.07.23.736688

**Authors:** Cravero Charlotte, Lesur-Kupin Isabelle, Choisne Nathalie, Leple Jean-Charles, Vassilieff Helena, de Miguel Marina, Gautier Véronique, Belmonte Elodie, Pailler Vincent, Poncet Charles, Dia Sow Mamadou, Huneau Cecile, Klopp Christophe, Ehrenmann Francois, Giovanni G Vendramin, Alía Ricardo, Bellec Arnaud, Panaud Olivier, Maumus Florian, Salse Jérôme, Pichot Christian, Plomion Christophe, Marande William

## Abstract

Conifers, which comprise nearly two-thirds of extant gymnosperm species, are ecologically and economically important but remain genomically understudied because of their exceptionally large, repeat-rich genomes. Here, we report a chromosome-level assembly of the haploid genome of *Cupressus sempervirens* generated using PacBio HiFi reads and scaffolded with optical and genetic maps. The 10 Gb assembly shows exceptional contiguity for a conifer genome (contig N50 = 29.8 Mb) and was organized into 11 pseudomolecules. Iso-Seq-supported annotation identified 42,980 protein-coding genes. Repetitive elements account for over 80% of the genome, with LTR retrotransposons alone representing 52.5%. Transposable elements (TE) are pervasive in both intergenic and genic regions and have a major impact on gene architecture: TE insertions within introns generate ultra-long introns, often exceeding 100 kb, and drive gene size expansion. Analyses of LTR retrotransposon dynamics indicate that genome enlargement in *C. sempervirens* was driven not by recent transpositional bursts, but by the long-term accumulation and incomplete removal of ancient LTR retrotransposons. Consistent with this pattern, paleogenomic reconstruction across representative gymnosperms found no evidence of whole-genome duplication in the Cupressus lineage. This reference genome provides a valuable resource for studying conifer genome evolution, gene structure, and traits of agronomic and ecological interest, including cypress pollinosis.

## Introduction

Conifers, among Earth’s most ancient plant lineages, dominate boreal and temperate forest ecosystems (Farjon and Filer 2013). With over 600 extant species, comprising two-thirds of all gymnosperms, they are vital for global biodiversity, carbon sequestration, and timber production. Their resilience across extreme environments, from deserts to mountains, reflects unique physiological adaptations: allogamous reproduction sustaining high genetic diversity (Leslie et al. 2018), (“Conifer Reproductive Biology,” n.d.) and specialized metabolic pathways such as terpene synthesis, which play a key role in biotic and abiotic stress responses (Loreto and Schnitzler 2010; Bohlmann and Keeling 2008; Celedon and Bohlmann 2019). These traits have enabled conifers to thrive for over 300 million years.

However, their complex genomes, often exceeding 20 Gbp with high heterozygosity, have long hindered sequencing efforts (“Plant DNA C-Values Database,” n.d.), despite their potential to inform tree improvement programs facing climate change (Mackay et al. 2012). Recent advances in sequencing technologies (Sun et al. 2022) have overcome some of these barriers, enabling high-quality assemblies for emblematic species such as Japanese cedar, Chinese pine, and giant sequoia (Scott et al. 2020; Song et al. 2021; Niu et al. 2022; Fujino et al. 2024).

The massive size of gymnosperm genomes (3 Gbp to over 30 Gbp) is largely driven by transposable elements (TEs), which constitute 69–88% of genomic content in species like *Pinus tabuliformis* and *Welwitschia mirabilis* (Scott et al. 2020; Song et al. 2021; Niu et al. 2022; Fujino et al. 2024; Wan et al. 2021). Notably, TEs extensively insert into genic regions, resulting in the formation of giant genes with ultra-long introns (>100 kb) (Scott et al. 2020; Song et al. 2021; Niu et al. 2022; Fujino et al. 2024). Two non-exclusive hypotheses have been proposed to explain this genomic “obesity”: the accumulation of TEs without efficient removal, and ancient whole-genome duplication (WGD) events (Wan et al. 2018 and 2021; Li et al. 2015). However, the paleohistory of this clade remains poorly understood.

Among European conifers, the Mediterranean cypress (*Cupressus sempervirens* L.) is a key species, prized since antiquity for its ornamental value, durable wood, and role as a windbreak (Weick et al. 2023). Native to the eastern Mediterranean and widely introduced globally (Caudullo and de Rigo 2016), *C. sempervirens* offers a unique opportunity for genomic analysis. Here, we leveraged on a haploid line derived from *Cupressus dupreziana via* male apomixis, (Caudullo and de Rigo 2016; Pichot et al. 2008) (Fig. 1a), which eliminates heterozygosity-related complexities and facilitates high-contiguity genome assembly. Using this resource, we aimed to (i) characterize the genic structure of *Cupressus*, focusing on TE impact in gene size, and (ii) elucidate the origins of its genome expansion through comparative analyses of TE dynamics and paleogenomics across gymnosperms.

**Figure 1:**
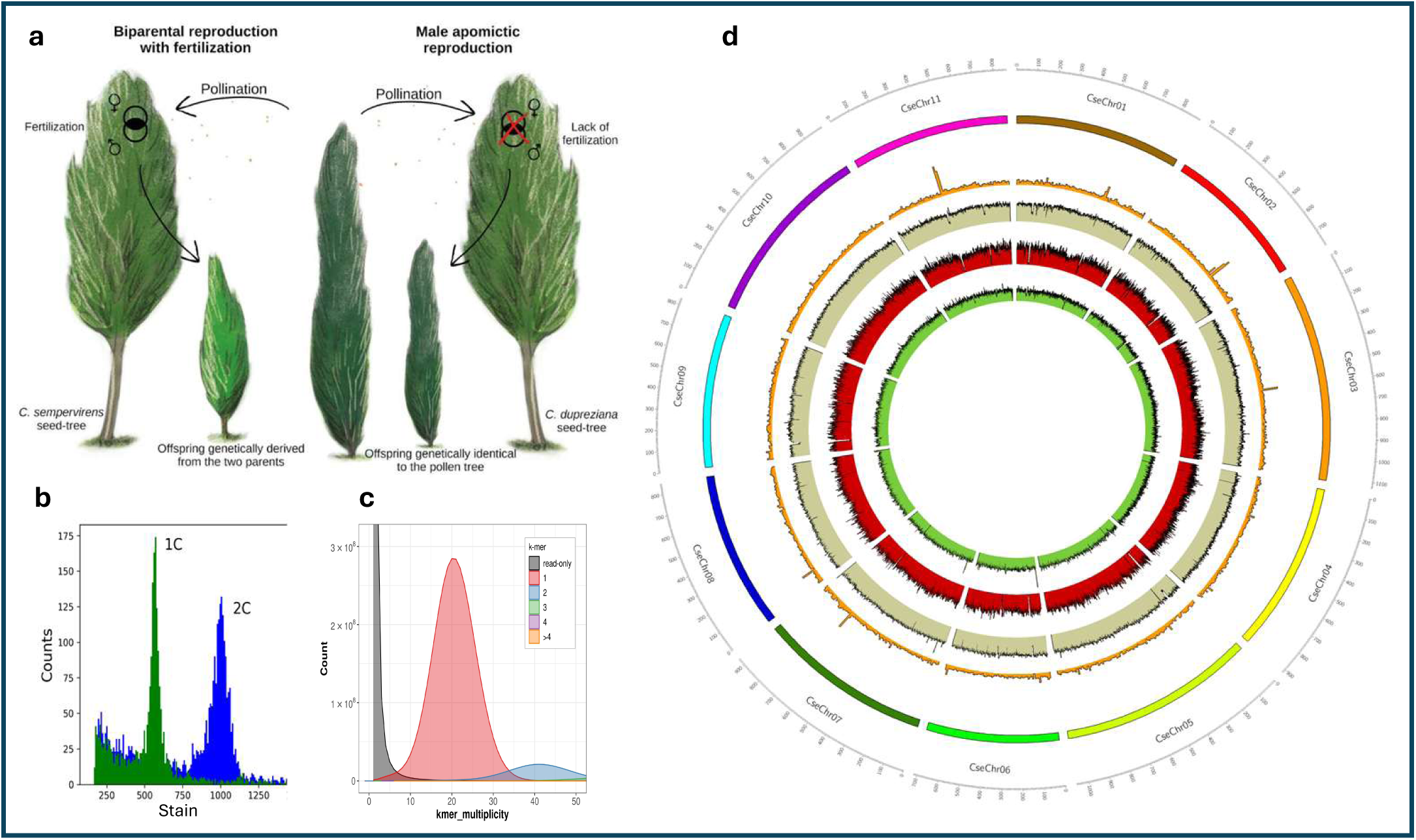
High quality *Cupressus sempervirens* genome assembly and annotation of an haploid line resulting from male apomictic reproduction using *C. dupreziana* as a surrogate seed-tree. a) Biparental reproduction in *Cupressus sempervirens* (left panel) leading to diploid offspring versus male apomictic reproduction using the *C. dupreziana* seed-tree (right panel) leading to haploid *C. sempervirens* offspring genetically identical to the C. sempervirens pollen tree, b) Flux cytometry of *C. sempervirens* haploid (green) and diploid (blue) lines, c) Merqury k-mer analysis of the genome assembly illustrating the lack of heterozygosity, d) Circos plot of genome feature : each track from outer to inner represents chromosomes, gene density, TE density, LTR copia density and LTR gypsy density.

## Results

### Genome assembly and scaffolding

From 215.7 Gb of PacBio HiFi reads generated using 10 Sequel II SMRT Cells, we assembled a 10.13 Gb haploid genome of *C. sempervirens* using Hifiasm. This assembly, in line with flow cytometry genome size prediction (Fig. 1b), comprised 1,549 contigs with an N50 of 29.8 Mb and a complete BUSCO score of 97.2%. Quality assessment via Merqury’s k-mer analysis confirmed its accuracy, yielding a quality score of 67 (equivalent to one error per 10 million base pairs) and 99.09% completeness between the reads and the assembly, supported by 21X coverage. The haploid state of the genome, critical for achieving high contiguity, was further validated by both flow cytometry and k-mer distribution (Fig. 1b and c).

For scaffolding, we generated 1.2 Tb of optical mapping data (>150 kb molecules), producing 251 optical maps (N50 = 215 Mb, 76x coverage) with 10.09 Gbp total size. Alignment of contigs to these maps yielded a 10 Gb scaffolded genome distributed across 36 scaffolds (N50 = 540 Mb) (Table 1). Finally, the alignment with a genetic linkage map of *Cryptomeria japonica* (Hasegawa et al. 2018), produced a chromosome-scale assembly of 11 pseudomolecules (717–1,126 Mb), matching the expected haploid chromosome number (total size: 9.98 Gb; Table 1, Fig. 1d). Telomere analysis confirmed that 8 pseudomolecules possess complete telomeric sequences at both ends (Fig. S1).

**Table 1:**
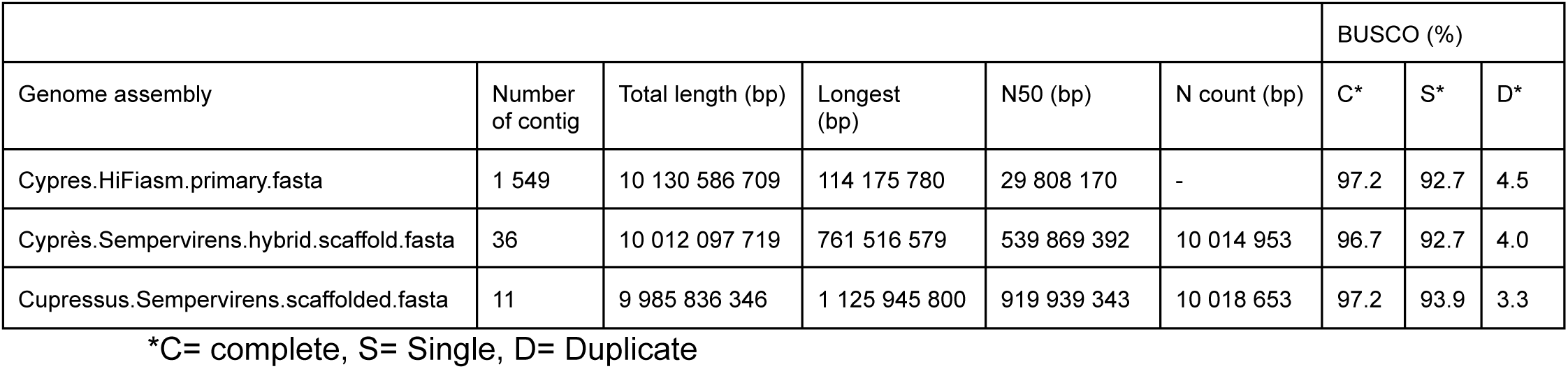
Genome assembly and scaffolding metrics.

|  |  |  |  |  |  | BUSCO (%) |  |  |
| --- | --- | --- | --- | --- | --- | --- | --- | --- |
| Genome assembly | Number of contig | Total length (bp) | Longest (bp) | N50 (bp) | N count (bp) | C* | S* | D* |
| Cypres.HiFiasm.primary.fasta | 1 549 | 10 130 586 709 | 114 175 780 | 29 808 170 | - | 97.2 | 92.7 | 4.5 |
| Cypres.Sempervirens.hybrid.scaffold.fasta | 36 | 10 012 097 719 | 761 516 579 | 539 869 392 | 10 014 953 | 96.7 | 92.7 | 4.0 |
| Cupressus.Sempervirens.scaffolded.fasta | 11 | 9 985 836 346 | 1 125 945 800 | 919 939 343 | 10 018 653 | 97.2 | 93.9 | 3.3 |

\*C= complete, S= Single, D= Duplicate

### Structural and functional gene annotation

Using the 150,650 IsoSeq sequences produced in this study, we annotated the *C. sempervirens* genome with the Eugene software (Carrere et al. 2023; Sallet et al. 2014). This analysis identified 77,322 genes, including 42,980 protein-coding gene models, with an average of 4 exons (mean size of 402 bp) and 3 introns (mean size of 6,202 bp) per gene and an average CDS length of 1,063 bp (Table 2). BUSCO assessment (v5.4.3, viridiplantae_odb10) confirmed assembly completeness, with 97.2% complete genes, while proteins reached 84.7% complete, 9.4% fragmented, and 5.9% missing models.

**Table 2:** Structural and functional annotation : main metrics.

| Number of genes | Number of protein coding genes (with functional annotation) | mean gene length (Kb) | mean CDS length (bp) | mean exon length (bp) | mean intron length (bp) | number of introns >10 kb |
| --- | --- | --- | --- | --- | --- | --- |
| 77,322 | 42,980 (39052) | 18,3 | 1,062 | 402 | 6,202 | 21,053 |

Protein validation and functional annotation revealed that 38,671 proteins (89.97%) were validated by Psauron (confidence score > 0.5), while 4,309 (10.03%) scored below 0.5, suggesting potential out-of-frame CDS. Functional annotations were assigned to 39,430 proteins (91.74%) via InterProScan (39,052; 90.86%), BLASTKoala (10,265; 23.88%), eggNOG (36,683; 85.35%), and E2P2 (20,764; 48.31% enzymatic functions). Only 3,550 proteins (8.26%) remained unannotated, of which 86.65% (3,076) were small (<100 amino acids) (Table 2; Fig. S2).

Additionally, we annotated long non-coding RNAs (lncRNAs). From 27,596 novel transcripts (identified via Cufflinks), 26,891 transcripts (20,987 genes) were classified as lncRNAs by FEELnc, with lengths ranging from 260 bp to 27,819 bp (mean: 4,077 bp).

### Transposable Elements (TE) content and annotation

To characterize the repetitive landscape of the *C. sempervirens* genome, we constructed a *de novo* repeat library using a TEdenovo pipeline (Flutre et al. 2011; Hoede et al. 2014) and annotated each chromosome with the TEannot pipeline (Quesneville et al. 2005) from the REPET package v3.0. Repetitive elements dominate the genome, representing 81.48% to 83.57% of each chromosome, with Class I TEs (retrotransposons) contributing to 57.8% of the assembly (Table 3; Table S3). These elements exhibit a uniform genomic distribution, with no detectable chromosomal bias (Fig. 1d).

**Table 3:** TE families and respective genome coverage.

| Class | TE | coverage (bp) | coverage/genome (%) |
| --- | --- | --- | --- |
| *Retrotransposons* | Copia | 2153382911 | 21,508 |
|  | Gypsy | 2613554858 | 26,104 |
|  | LARD | 430675790 | 4,302 |
|  | TRIM | 21563483 | 0,215 |
|  | Retrotransposon | 35880676 | 0,358 |
|  | Penelope | 12909227 | 0,129 |
|  | LINE-L1 | 498360912 | 4,978 |
|  | LINE-RTE | 19152884 | 0,191 |
|  | SINE | 590045 | 0,006 |
| *DNA transposons* | Sola2 | 180044961 | 1,798 |
|  | CACTA | 133861465 | 1,337 |
|  | hAT | 51220996 | 0,512 |
|  | PIF-Harbinger | 3279908 | 0,033 |
|  | MuDR | 530531 | 0,005 |
|  | MITE | 860729 | 0,009 |
|  | TIR | 2058061 | 0,021 |
|  | TIR-nonautonomous | 72605372 | 0,725 |
|  | Helitron | 40423932 | 0,404 |
| Confused TE | Confused | 1330290640 | 13,287 |
| Unclassified sequence | Unclassified | 815067932 | 8,141 |
| Potential Host gene | PHG | 22192150 | 0,222 |
| SSR | SSR | 56786562 | 0,567 |
| rDNA | rDNA | 8074012 | 0,081 |

Among Class I TEs, Long Terminal Repeat retrotransposons (LTR-RTs) are the most abundant, comprising 52.5% of the genome. The Gypsy superfamily (26.1%) predominates over Copia (21.5%), consistent with patterns observed in other conifers (Table 3). Non-LTR retrotransposons (LINE and SINE) account for 5.2% of the genome, with LINE-L1 as the predominant superfamily (4.9%), mirroring findings in *C. japonica (Fujino et al. 2024)*. Class II elements (DNA transposons) represent 4.8% of the genome, with Sola2 as the most abundant superfamily (Table 3). Unclassified TEs and repeats with ambiguous classification (named Confused TE) constitute 8.1% and 13.3% of the genome, respectively (Table 3).

Manual curation of the repeat library identified additional elements, including 0.2% host genes, 0.5% SSR sequences, and 0.08% rDNA sequences. We also detected Penelope superfamily retrotransposons (0.1%), previously reported in *Pinus taeda* and *Picea abies* (Table 3). Notably, endogenous caulimovirid elements (ECVs), widespread in angiosperms and recently documented in gymnosperms (Diop et al. 2018), cover ∼36 Mbp (0.36%) of the *C. sempervirens* genome (Fig. S3). Phylogenetic analysis indicated that *C. sempervirens* reverse-transcriptase sequences are distributed across two distantly related groups, both of which are significantly divergent from reference caulimovirid sequences and hence probably represent novel genera (Fig. S4).

### Gene size and ultra-long intron

To investigate the structural composition of genic regions in C. sempervirens, we analyzed exons and introns, which collectively span 786.6 Mb (7.9% of the genome), while coding sequences (CDS) account for only 65 Mb (0.65%). This disparity underscores the dominance of intronic sequences in conifer gene architecture. Adjusting the maximum intron length parameter from 6 kb to 150 kb during structural annotation dramatically improved the BUSCO completeness score (from 40% to 85%), confirming the presence of ultra-long introns (>100 kb) (Fig. 2), as previously described in gymnosperms (Fujino et al. 2024; Niu et al. 2022).

**Figure 2:**
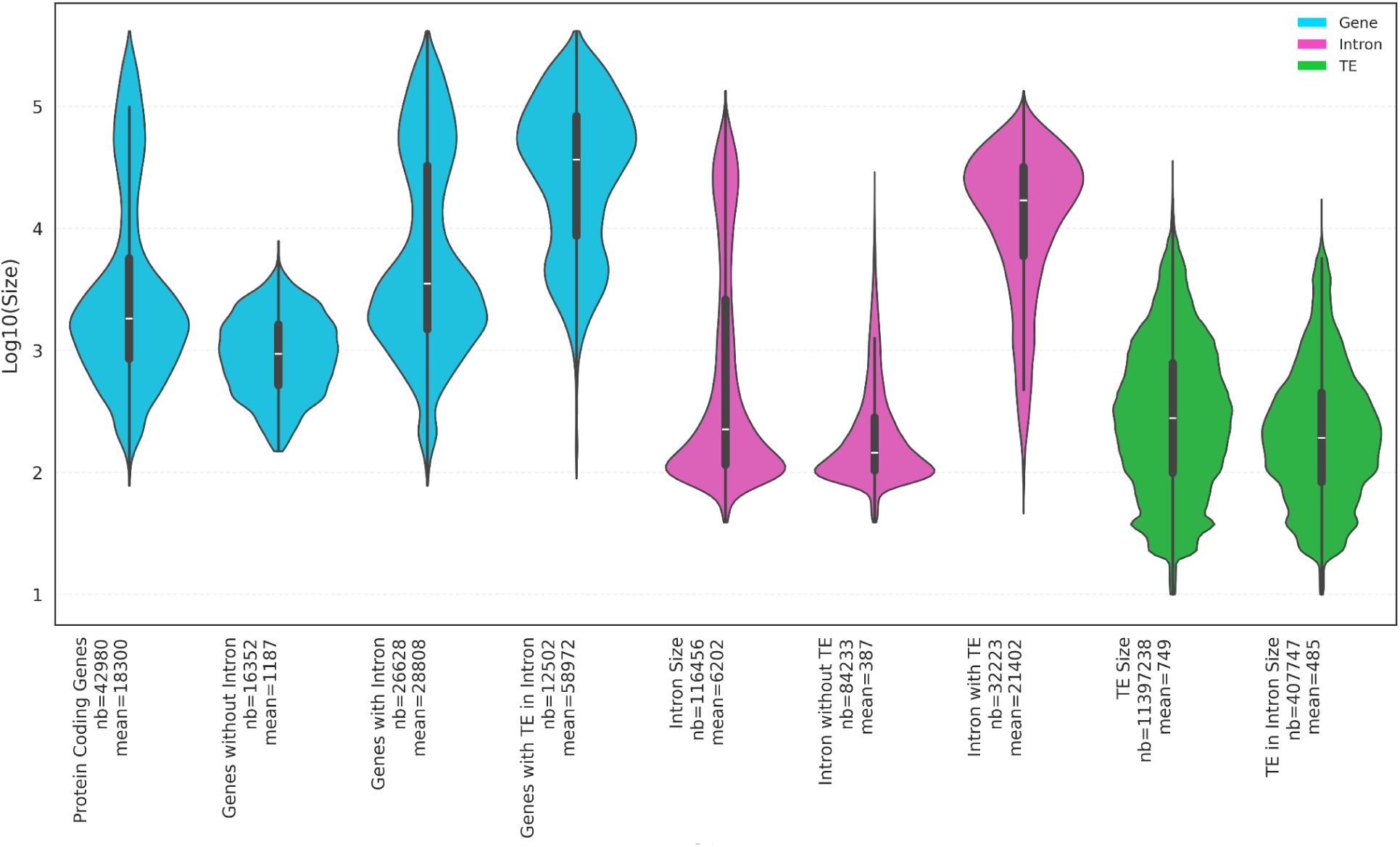
Size distribution of genes, introns and TE. Violin plots showing the log (10) size distribution and impact of TE insertion in genes or introns. Each violin plot shows a miniature box-and-whisker plot giving the median (horizontal line), the upper and lower quartiles (rectangular box), and the extent of gene length distribution (whiskers). The average length for each feature is shown on the Y axis.

We hypothesized that TE insertions contribute to these exceptionally large introns. Using the TE annotation, we identified 32,454 TE-containing introns out of ∼116,000 total introns, affecting 18,224 of the 42,980 annotated genes. The size distributions of genic features, visualized in violin plots (Fig. 2), reveal striking patterns:

- Protein-coding genes (blue) exhibit a bimodal size distribution (mean = 18,300 bp), where the first peak corresponds to intronless genes (mean = 1,187 bp), and the second peak reflects genes with introns (mean = 28,808 bp). The long tail of this distribution, extending to genes >400 kb, is driven by TE-containing introns (mean = 58,972 bp), which inflate gene size.
- Introns (pink) show a right-skewed distribution, with TE-free introns (mean = 387 bp) sharply peaking at smaller sizes, while TE-containing introns (mean = 21,402 bp) display a broader distribution, reflecting the presence of ultra-long introns.
- TEs (green) exhibit a log-normal-like distribution, with intron-associated TEs (mean = 487 bp) being smaller than global TEs (mean = 749 bp), suggesting selective compaction of TEs within introns.

These distributions demonstrate that TE insertions within introns are the primary driver of gene size expansion in *C. sempervirens*, reshaping its genomic landscape.

### Genome size investigation

#### Evolutionary Dynamics of LTR Retrotransposons and Genome Size Expansion

The TE annotation of the *C. sempervirens* genome reveals that Class I elements (retrotransposons) dominate the non-genic compartment, contributing 5.8 Gbp—a pattern consistent with other gymnosperms (e.g., *Picea abies*, *Pinus taeda*) and large angiosperm genomes (Russo et al. 2024; Liang et al. 2025; Wicker et al. 2018; Oliver et al. 2013). While angiosperms efficiently eliminate TE-related sequences via intra- or inter-element recombination (Tian et al. 2009; Vitte et al. 2007), the persistent expansion of gymnosperm genomes—including *Cupressus*—suggests differential TE turnover dynamics. To test whether this expansion reflects recent transpositional bursts or reduced TE elimination, we compared LTR retrotransposon dynamics between *Cupressus* (10 Gb) and *Zea mays* (2 Gb), a model for TE turnover in angiosperms (Tian et al. 2009; Vitte et al. 2007; El Baidouri and Panaud 2013). Contrary to the “recent bursts” hypothesis, we identified 49,019 intact LTR retrotransposons in *Cupressus*, fewer than in maize (66,075), despite *Cupressus*’ fivefold larger genome. Insertion age estimates further revealed that retrotranspositional activity in *Cupressus* is not only less intense but also older than in maize (Fig. S5). These findings support a model where *Cupressus*’ massive genome size stems from ancient, poorly eliminated TE bursts, rather than recent proliferation.

To explore the mechanisms underlying this TE retention, we focused on unequal recombination (UR), a key pathway for TE removal via solo-LTR formation (Tian et al. 2009; Vitte et al. 2007). While *Pinaceae* (e.g., *Picea abies*) exhibit low UR rates (solo-to-intact LTR ratio ∼1:9), our analysis in *Cupressus*, using LTR retriever, yielded a solo-to-intact LTR ratio of 2.8:1, indicating that UR occurs but remains insufficient to counteract TE accumulation. This reduced TE purging efficiency may reflect altered recombination pathways in *Cupressaceae*, contributing to the long-term retention of ancient LTRs and the expansion of intergenic spaces.

#### Paleogenomic Reconstruction of Conifer Genome Evolution

Beyond TE proliferation, polyploidization represents another potential driver of genome size expansion in gymnosperms. To elucidate the paleohistory of the gymnosperm-conifer complex and identify past polyploidization events, we performed a comparative genomic analysis of eight representative species: *Sequoiadendron giganteum* (Sg), *Cupressus sempervirens* (Cu), *Taxus chinensis* (Tch), *Pinus tabuliformis* (Pit), *Ginkgo biloba* (Gb), *Cycas panzhihuaensis* (Cy), *Welwitschia mirabilis* (Ww), and *Gnetum montanum* (Gn), using *Amborella trichopoda* (Amt) as an outgroup. Alignments and ancestral genome reconstructions followed methods described previously (Pont et al. 2019; Murat et al. 2017; Siguret et al. 2026) (Fig. 3a).

**Figure 3:**
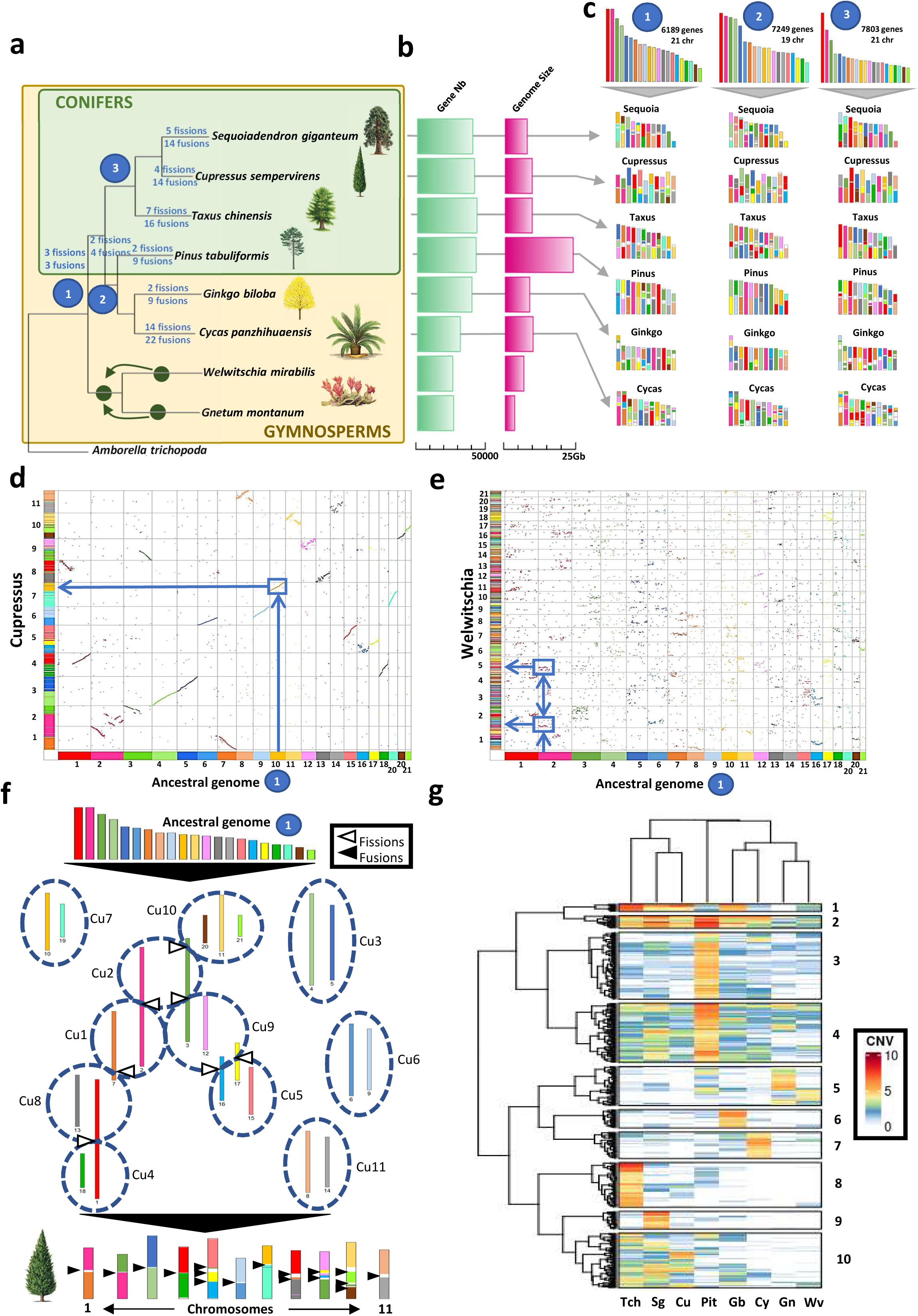
Conifers’ paleoevolutionary history. **a.** Evolutionary scenario of the modern *Sequoia* (Sg), *Cupressus* (Cu), *Taxus* (Tch), *Pinus* (Pit), *Ginkgo* (Gb), *Cycas* (Cy), *Welwitschia* (Ww), and *Gnetum* (Gn) genomes, with *Amborella trichopoda* (Amt) as an outgroup, from inferred ancestral karyotypes (1, 2 and 3), with duplication (WGD) events shown with green dots on the tree branches, along with shuffling events (fusions and fissions). **b.** Illustration of the large diversity in genomes structures, exemplified with the number of genes and the genome sizes. **c.** The modern genomes are illustrated with different colours reflecting their origins from the ancestral chromosomes of the three inferred ancestors (1, 2 and 3). **d and e.** Complete dot-plot based deconvolution of the observed paralogy and orthology (dot-plot diagonals) between the inferred Gymnosperm-conifer ancestral (the colors on the axis illustrates the ancestral chromosomes), *Cupressus* (with no sign of duplication with a one-to-one chromosome relationship, d) and *Welwitschia* (with sign of duplication with a one-to-two chromosome relationship, e) genomes **f.** Evolutionary scenario of rearrangements of ancestral chromosomes in shaping the modern *Cupressus* genome through 17 fusions (black arrows) and 7 fissions (white arrows) **g.** Clustering of 551 genes (lines) showing Copy Number Variation (CNV, see color legend at the right) between the investigated species (in columns), defining 10 groups of genes with different contrasts in CNVs between subsets of species.

Despite diverse modern genome structures (Fig. 3b), conserved synteny analyses allowed us to infer three ancestral karyotypes (Table S4)

- Sequoia-Cupressus-Taxus ancestor: 7,803 genes, 21 chromosomes.
- Pinus-Ginkgo-Cycas ancestor: 7,249 genes, 19 chromosomes.
- Basal gymnosperm ancestor (Sequoia-Cupressus-Taxus-Pinus-Ginkgo-Cycas): 6,189 genes, 21 chromosomes (Fig. 3c).

These reconstructions largely refine previous models of conifer karyotype evolution, sketched from comparative linkage map analysis (de Miguel et al., 2015), which proposed that modern genomes (typically 11–12 chromosomes) derived from 19–21 ancestral chromosomes *via* ancestral chromosome fusion and fission events. Our comparative analysis reveals a 1:1–2 ancestral-to-modern chromosome correspondence, highlighting lineage-specific polyploidization events. Notably, *Welwitschia* and *Gnetum* exhibit clear evidence of a recent, possibly shared WGD, while *Cupressus* shows no WGD signature (Fig. 3d–e). This rules out WGD as a contributor to *Cupressus*’ massive genome size, reinforcing the role of TE accumulation as the primary driver of expansion.

Finally, we modeled the evolutionary trajectory of modern genomes from the inferred gymnosperm ancestor karyotypes using a parsimony-based approach (minimizing chromosomal shuffling, Murat et al. 2017; Siguret et al. 2026). This analysis proposes that the *Cupressus* genome evolved through 17 fusions and 7 fissions (Fig. 3f), maintaining a structure closer to the ancestral state than other conifers.

#### Gene Family Expansion: Functional Specialization in Conifers

To explore the functional diversification within the gymnosperm-conifer complex, we analyzed copy number variations (CNVs) among 551 conserved genes across the eight studied species. These genes cluster into 10 distinct groups based on their CNV patterns, revealing lineage-specific expansions that may underpin ecological and evolutionary adaptations:

- Group 1 (12 genes): High copy number in *Tch*, *Sg*, *Cu*, *Gb*.
- Group 2 (20 genes): Low copy number in *Gn*, *Ww*.
- Group 3 (106 genes): High copy number in *Pit*.
- Group 4 (94 genes): Variable copy numbers across all species.
- Group 5 (62 genes): High copy number in *Gn*, *Ww*.
- Group 6 (29 genes): High copy number in *Gb*.
- Group 7 (42 genes): High copy number in *Cy*.
- Group 8 (71 genes): High copy number in *Tch*.
- Group 9 (28 genes): High copy number in *Sg*.
- Group 10 (87 genes): High copy number in *Tch*, *Sg*, *Cu*, *Pit*, *Gb*.

These differential CNV patterns suggest that gene family expansions may have played a key role in the emergence of species-specific traits. Notably, the absence of recurrent polyploidization events, as previously reported in conifers (Scott et al., 2020; Xiong et al., 2021; Niu et al., 2022; Liu et al., 2022), highlights gene duplication and diversification as the primary mechanisms driving functional innovation in this lineage. This aligns with our earlier findings on TE-driven genome expansion and chromosomal stability, reinforcing the idea that conifers rely on gradual genomic changes rather than abrupt polyploidization events to adapt and diversify.

## Discussion

Gymnosperm genomes have long posed major assembly challenges because of their large size, high repetitive content, and, in many species, substantial heterozygosity (Stevens et al. 2016). Recent long-read sequencing technologies, particularly PacBio HiFi, have markedly improved assembly continuity in conifers, as illustrated by chromosome-scale genomes such as *C. japonica* (Fujino et al. 2024). In this context, our haploid-derived assembly of *C. sempervirens* achieves the highest contiguity amongst conifer genome, with a contig N50 of 29.9 Mb. The use of a homozygous line substantially reduced the complications associated with heterozygosity, and integration with optical and genetic maps enabled chromosome-scale scaffolding, including eight telomere-to-telomere chromosomes. Beyond its technical quality, this assembly provides a useful framework for interpreting the evolutionary forces that have shaped *Cupressaceae* genomes and gymnosperm genome architecture more broadly.

The approximately 10-Gb genome of *C. sempervirens*, although at the lower end of the gymnosperm size range (Pellicer and Leitch 2020), is strongly dominated by repetitive DNA. Class I TE account for 5.8 Gb, representing 57.8% of the genome, with LTR retrotransposons contributing the largest fraction. Remarkably, this TE-rich architecture extends beyond intergenic space and also affects gene structure, as many introns contain abundant TE-derived sequences and contribute to highly expanded gene bodies. This pattern is consistent with one of the defining features of conifer genomes, in which large introns are a major component of genome expansion rather than a marginal feature (Nystedt et al. 2013; Niu et al. 2022; Song et al. 2021; Fujino et al. 2024). Importantly, these enlarged gene structures do not appear to impose a simple transcriptional penalty. In *P. tabuliformis*, for example, large genes with very long introns can remain highly expressed, suggesting that conifer genomes have evolved regulatory and splicing systems compatible with highly expanded gene bodies (Nystedt et al. 2013; Niu et al. 2022; Song et al. 2021). Long introns should therefore not be viewed merely as passive by-products of genome enlargement, but as stable components of conifer genome architecture that may influence gene regulation, chromatin structure, and genome organization.

More broadly, the repeat-rich composition of the *C. sempervirens* genome is consistent with the prevailing view that conifer genome enlargement is driven mainly by long-term transposable-element accumulation rather than by unusually high gene number or recent whole-genome duplication (Nystedt et al. 2013; Niu et al. 2022; Song et al. 2021). In angiosperms such as maize, TE landscapes are often shaped by relatively rapid turnover through unequal recombination and related deletion mechanisms (Tian et al. 2009; Vitte et al. 2007; El Baidouri and Panaud 2013). By contrast, conifer genomes appear to retain many TE insertions over much longer evolutionary timescales. However, the solo-LTR:intact-LTR ratio observed in *C. sempervirens* suggests that repeat removal in this species may be more active than in *Picea abies*, while still remaining incomplete relative to many angiosperms. Rather than supporting a single conifer-wide model of uniformly inefficient TE elimination, our results point to variation in repeat-turnover dynamics among gymnosperm lineages, consistent with recent comparative studies indicating that *Pinaceae* and *Cupressaceae* differ in the balance between TE accumulation and removal (Nystedt et al. 2013; Niu et al. 2022; Song et al. 2021; Fujino et al. 2024; Wan et al. 2022).

Several factors may contribute to the persistence of TEs in *C. sempervirens* and other conifers. Long generation times and reduced effective recombination may limit the efficiency of recombination-based repeat removal (Leitch and Leitch 2012; Niu et al. 2022), while epigenetic silencing mechanisms may stabilize TE-rich regions without physically eliminating them. In *P. tabuliformis*, high levels of DNA methylation, especially CHG methylation, have been proposed to play a dual role in repeat silencing and in maintaining transcriptional function across large, repeat-rich genes (Nystedt et al. 2013; Niu et al. 2022; Song et al. 2021). It is therefore plausible that similar mechanisms contribute to the long-term retention of ancient TE insertions in *Cupressaceae*. At the same time, whether retained TEs also provide a significant reservoir of regulatory or adaptive variation in conifers remains largely unresolved and will require broader comparative and functional analyses.

Our paleogenomic reconstruction further suggests that the large size of the *C. sempervirens* genome cannot be explained by ancestral WGD. Instead, the modern genome appears to have arisen through a history of chromosome rearrangements, including fusions and fissions, together with extensive repeat accumulation and local duplication. This interpretation is broadly consistent with assembly- and synteny-based studies of conifer genomes, which generally do not recover evidence that recent or moderately ancient WGD events are the primary drivers of conifer genome gigantism (Nystedt et al. 2013; Niu et al. 2022; Song et al. 2021; Zhang et al. 2025). Rather, these studies support a model in which genomic innovation in conifers is associated with small-scale duplication, copy-number variation, and lineage-specific gene family expansion, especially in pathways related to stress response and defense, while intergenic and intronic expansion accounts for much of the increase in genome size (Nystedt et al. 2013; Niu et al. 2022; Song et al. 2021). In this sense, conifer genomes appear to have achieved substantial genomic plasticity without relying on recurrent polyploidization, in marked contrast to the pattern commonly invoked for angiosperm evolution (Soltis et al. 2015).

Nevertheless, the role of ancient WGD in gymnosperm evolution remains debated. Some phylogenomic analyses have inferred older duplication events in seed plants, gymnosperms, or major conifer subclades (Li et al. 2015; Celedon and Bohlmann 2019; Liu et al. 2021), and these conflicting results indicate that the paleopolyploid history of gymnosperms is not fully resolved. In our analyses, clear WGD signatures were detected only in the lineages leading to *Welwitschia mirabilis* and *Gnetum montanum*, suggesting that polyploidization may have been lineage-specific rather than a shared feature of gymnosperm evolution. Whether this pattern reflects ecological selection, lineage-specific differences in duplicate retention, or other aspects of genome biology remains unclear. Notably, even in lineages where WGD is inferred, TE dynamics still appear to play an important role, reinforcing the view that gymnosperm genome evolution is shaped by varying combinations of duplication, repeat proliferation, and structural reorganization rather than by a single universal model (Li et al. 2015; Wan et al. 2021).

Finally, beyond its evolutionary significance, the *C. sempervirens* genome provides a valuable resource for cypress genetics and breeding. In addition to traits such as growth form, ornamental value, and tolerance to pathogens or environmental stress, pollen production is an important target in *Cupressus* and more broadly in *Cupressaceae* because cypress pollen is a major cause of seasonal respiratory allergy in many regions. A high-quality reference genome should facilitate the identification of genes and regulatory pathways involved in male reproduction and allergen production, thereby supporting both basic research and breeding strategies aimed at developing low-pollen or male-sterile varieties. In this regard, the production of haploid lines through the surrogate capacity of *C. dupreziana*, such as the ‘cs-2001-9-10’ sequenced in this study, offers another and direct alternative for the production of low- or non-pollinating varieties. Indeed, haploid status disrupts normal microsporogenesis and leads to strongly altered pollen production. On this basis, ‘cs-2001-9-10’ is also a newly patented cypress variety, called ‘Sirocco’, selected for the prevention of pollinosis. It is now in the process of vegetative propagation for commercialisation in the coming years.

The genomic characterization of such material therefore has both biological and applied relevance, and provides a foundation for integrating reproductive biology, genome evolution, and cultivar development in cypress.

## MATERIAL and METHODS

### Plant material

The unusual haploid line of *C. sempervirens* targeted in this project was generated using *Cupressus dupreziana* as a surrogate mother. *C. dupreziana* is *a* highly endangered Saharan cypress from the Tassili N’Ajjer plateau and it has a unique reproductive system based on male apomixis (Pichot et al 2001). Other unusual aspects of reproductive biology in Mediterranean cypresses that contributed to the creation of this unique genetic resource include tetraspory (El Maâtaoui et al 1998) and endosperm polyploidy (Pichot & El Maataoui 1997). Sexual reproduction in *C. dupreziana* is the only known example of male apomixis in plants, which consists in the process of development of an embryo from unreduced diploid pollen, in the maternal tissues that do not contribute to the genetic material of this embryo. Surprisingly, *C. dupreziana* trees also act as a surrogate mother for other species, allowing the development of progeny from reduced haploid pollen. In this study, pollination of *C. dupreziana* female cones with *C. sempervirens* pollen led to the production of haploid *C. sempervirens* genotypes trees (Pichot et al 2008).

All plant material used for DNA and RNA extraction were collected from an 11-year-old cypress tree as a ramet produced by vegetative propagation of clone ‘cs-2001-9-10’. This clone was selected among hundreds of haploid lines, based on growth and shape characteristics, with the objective of providing a new variety for cypress pollinosis prevention.

### Ploidy level determination by flux cytometry

Several independent haploid lines have been produced using this method by Christian Pichot (INRAE PACA, Avignon, France), and were available for this project.

Ploidy was evaluated by flow cytometry by comparing samples collected from the studied tree to that of diploid cypress as control. Somatic tissues were chopped with a razor blade, filtered through a 30-µm CellTricsTM filter and stained in 1.5 ml of solution B (4′,6-diamidino-2-phenylindole, DAPI staining) of the High resolution DNA kit, type T (PARTEC, Münster, Germany). The DNA content of the nuclei was evaluated using a PARTEC Ploidy Analyser PA equipped with a HBO lamp for UV excitation. Measurements were assigned according to relative fluorescence intensity. Results were stored in histogram-type data files and studied with the WINMDI software (version 2.7, copyright© 93–98 Joseph Trotter).

### Plant tissues sampled

Samples were collected on 13 December 2021 from 5 different tissues: scale leaves, young stem, mature xylem, phloem and immature strobiles. All samples were flash frozen in liquid nitrogen and then stored at −80 °C.

### HMW DNA extraction, SMRTbell Libraries preparation and sequencing

High molecular weight (HMW) DNA was extracted from 2 g of needle using QIAGEN Genomic-tips 500/G kit (Qiagen, MD, USA) following the manufacturer’s protocol for tissue extraction. Briefly, 2 g of young needle tips material were grinded in liquid nitrogen with mortar and pestle. After 3 hours of lysis and one centrifugation step, the DNA was immobilized on the column. After several washing steps, DNA was eluted from the column, then desalted and concentrated by Isopropyl alcohol precipitation. A final wash in 70% ethanol was performed before resuspending the DNA in EB buffer.

Analyses of DNA quantity and quality were performed using NanoDrop and Qubit (Thermo Fisher Scientific, MA, USA). DNA integrity was also assessed using the Agilent FP-1002 Genomic DNA 165 kb kit on the Femto Pulse system (Agilent, CA, USA).

HiFi libraries were produced using SMRTbell Express 2 Template prep kit, following the “procedure and checklist - preparing HiFi SMRTBell Libraries using SMRTbell Express Template prep kit 2.0” protocol.

High Molecular Weight Genomic DNA (15 ug) was sheared with the 25 kb program using a Diagenode Megaruptor (Diagenode) generating DNA fragments of approximately 20 kb. A Femto Pulse (Agilent Technologies, Santa Clara, CA, USA) assay was used to assess the fragments size distribution. Sheared genomic DNA was used in the enzymatic reactions to remove the single-strand overhangs and to repair any damage that may be present on the DNA backbone. A A-tailing reaction followed by the overhang adapter ligation was conducted to generate the SMRTBell template. After nuclease treatment and a 1 X AMPure PB beads purification, the sample was size-selected using the BluePippin (Sage Science, Beverly, MA, USA) in order to recover all the material above 10 kb. The sample was then purified with 1 X AMPure PB Beads to obtain the final libraries around 20 kb. The SMRTBell library was quality inspected and quantified on a Femto Pulse (Agilent Technologies) and a Qubit fluorimeter with Qubit dsDNA HS reagent Assay kit (Life Technologies). A ready-to-sequence SMRTBell Polymerase Complex was created using a Sequel II Binding Kit 2.2 and the primer v5. The PacBio Sequel II instrument was programmed to load a 90 pM library and sequenced the sample with Sequel II Sequencing plate v2.0 (Pacific Biosciences), acquiring 2 hours of pre-extension and one movie of 30 hours per SMRTcell, using CCS mode.

### uHMW DNA extraction, optical map production and hybrid scaffolding

uHMW DNA was purified from 0.6 g of fresh young needle tips according to the Bionano Prep Plant Tissue DNA Isolation Base Protocol (30068 - Bionano Genomics) with the following specifications and modifications. Briefly, the needles were fixed in buffer containing formaldehyde. Following 3 washes, leaves were cut in 2 mm pieces and disrupted with rotor stator in the homogenization buffer containing spermine, spermidine and beta-mercaptoethanol. Nuclei were washed, purified using a density gradient and then embedded in agarose plugs. After overnight proteinase K digestion (Qiagen) in the presence of Lysis Buffer and one hour treatment with RNAse A (Qiagen), plugs were washed and solubilized with 2 µL of 0.5 U/µL AGARase enzyme (ThermoFisher Scientific). A dialysis step was performed in TE Buffer (ThermoFisher Scientific) to purify DNA from any residues. The DNA samples were quantified by using the Qubit dsDNA BR Assay (Invitrogen). The presence of mega base size DNA was visualized by pulsed field gel electrophoresis (PFGE).

Labelling and staining of the uHMW DNA were performed according to the Direct Label and Stain (DLS) protocol (30206 - Bionano Genomics). Briefly, labelling was performed by incubating 750 ng of genomic DNA with 1× DLE-1 Enzyme for 2 hours in the presence of 1× DL-Green and 1× DLE-1 Buffer. Following proteinase K digestion and DL-Green clean-up, the DNA backbone was stained by mixing the labelled DNA with DNA Stain solution in the presence of 1×Flow Buffer and 1× DTT, and incubating overnight at room temperature. The DLS DNA concentration was measured with the Qubit dsDNA HS Assay (Invitrogen, Carlsbad, CA, USA).

Labelled and stained DNA was loaded on one Saphyr chip and was run on the BNG Saphyr System according to the Saphyr System User Guide. Digitalized labelled DNA molecules were assembled to optical maps using the BNG Access software (solve version 3.6).

### RNA extraction, library preparation and sequencing

Prior to RNA extraction, the samples were finely milled with pestle and mortar in liquid nitrogen. About 100 mg of milled tissue was used to isolate separately total RNA from scale leaves, immature strobiles, mature xylem, phloem and young stems with a modified Chiang’s procedure (Chang et al. 1993). Briefly, after LiCl precipitation the RNA pellet was resuspended in 100 µl water and purified with RNeasy Plant Kit (Qiagen, France), according to manufacturer’s recommendations. On-column treatment with DNAse1 (Qiagen, France) ensures the elimination of genomic DNA during this purification step. RNA was eluted in DNAse-free water and quantified with a nanodrop spectrophotometer.

The Iso-Seq SMRTbell libraries were constructed according to the standard isoform sequencing protocol « Procedure and Checklist- Iso-Seq ™ Express Template Preparation for Sequel® and Sequel II Systems » using the NEBNext Single Cell/Low Input cDNA Synthesis & Amplification Module (New England Biolabs) and the ProNex Size-Selective Purification System (Promega) for size selection.

Each sample (300 ng total RNA) was first barcoded and then subjected to cDNA amplification using 12 cycles. Purified cDNAs were pooled in equal molarity for library preparation using the SMRTbell Express Template Prep Kit 2.0 (Pacific Biosciences) following the Iso-Seq protocol previously referenced. The library was prepared for sequencing by annealing primer v4 with the Sequel II Binding Kit 2.1. The PacBio Sequel II instrument was programmed to load a 80 pM library and sequenced the sample with Sequel II Sequencing plate v2.0 (Pacific Biosciences), acquiring one movie of 24 hours per SMRTcell.

### Isoseq sequence analysis

A total of 417,602 - 709,891 - 684,092 - 665,747 and 665,896 full-length non-concatemer transcript sequences were obtained for the five libraries prepared from scale leaves, immature strobili, mature xylem, phloem and young stem tissues, respectively. Combining these sequences, 317,881 high-quality consensus transcript sequences were obtained through Isoseq3-cluster analysis. A total of 311,250 transcripts were mapped to the genome sequence and further collapsed into a final set of 150,675 transcript sequences. Among those, we selected 150,650 transcript sequences larger than 200 bp.

### Genome assembly

Production of long-read sequences consisted of 10 SMRTCell proceeding in CCS mode. These sequences were corrected using the pbcromwell software embedded in SMRTLink (v10.1), the workflow manager for Sequel systems of PacBio, with the default settings (number of passes equal to 3). The corrected data were assembled using the HiFiasm assembler v0.15.5, (Cheng et al. 2021), thus producing a primary assembly and an alternative assembly (incomplete alternative assembly consisting of haplotigs under heterozygous regions). Assembly statistics were obtained using QUAST v5.0.2, (Gurevich et al. 2013), a quality assessment tool for genome assemblies. To assess the completeness of the genome assembly, we performed a BUSCO analysis with the viridiplantae database, v5.4.3 (Simão et al. 2015). In addition, we perform a k-mer analysis to check the quality of the dataset and the quality and completeness of the assembly using Merqury software (Rhie et al. 2020).

### Hybrid scaffolding between genome assembly and optical maps

A classic hybrid scaffolding between the sequence assembly and the optical genome maps was performed using the hybrid Scaffold pipeline from Bionano (https://bionano.com/wp-content/uploads/2023/01/30073-Bionano-Solve-Theory-of-Operation-Hybrid-Scaffold.pdf; solve version 3.6). Given the fact that the software couldn’t finish the hybrid scaffold, we splitted the analysis into several independent processes. First, we aligned the genome and the optical maps using the runChar 3.4 tool (https://bionano.com/software-downloads/). Then, we grouped all the contigs and optical maps matching together, thus creating 24 distinct datasets. We finally performed 24 independent hybrid scaffolding using the solve tool v3.6. This process allowed the achievement of the global hybrid scaffolding including all the contigs and all the optical maps.

### Anchoring to the genetic map of *Cryptomeria japonica*

Following the hybrid assembly, the consensus genetic map of Cryptomeria japonica (Hasegawa et al. 2018) was considered all together with the Allmaps software (kentUtils-v370, (Tang et al. 2015) for chromosome anchoring. A few incongruences between the genetic map and the Cupressus hybrid scaffolds were observed for linking groups 2, 5, 6 and 9. Scaffolds were broken in order to reconcile both data. We considered the consensus genetic map resulting from 4 different experiments of Cryptomeria species. The markers showed strong alignments with the closest relative species possessing a high-quality assembled genome, specifically the sequoia tree (Scott et al. 2020). Several manual rearrangements were performed according to the Cupressus sempervirens genetic map. Finally, telomeres were identified using FindTelomere (https://github.com/JanaSperschneider/FindTelomeres) and tidk (Brown et al. 2025).

### Repeats annotation

Due to the very large size of the Cupressus genome (10Gb), we developed a specific strategy to detect and annotate its repetitive elements.

We first used only the first 3 chromosomes (chr01, chr02 and chr03) to build a library of repetitive elements using a process implemented for large genomes to overcome computational challenges of TE discovery (Jamilloux et al. 2016).

The TEdenovo pipeline (Flutre et al. 2011; Hoede et al. 2014) from the REPET package v3.0 (https://urgi.versailles.inra.fr/Tools/) was used to build a library of consensus sequences representative of repetitive elements found in chr01, chr02 and chr03. To maximize repeat sampling across the whole genome, TEdenovo was launched independently on three 300 Mb subsets of effective sequence, each taken from one of the first three chromosomes. For each subset, sequences were cut at the level of stretches of >11 undefined bases (Ns) to generate « virtual » contigs.

A minimum of 5 sequences per cluster was forced to generate consensus sequences. Each sequence was classified using PASTEC (Hoede et al. 2014), followed by manual curation to identify and rename consensus sequences with ambiguous classification. After filtering out simple sequence repeats and unclassified sequences built with <10 copies per cluster, three libraries of 4,947, 4,522, and 4,664 consensus sequences were obtained for chr01, chr02 and chr03, respectively.

The TEannot pipeline (Quesneville et al. 2005) from the REPET package v3.0 was used with default parameters to independently annotate TE copies in each chr01, chr02 and chr03 using their respective library. To refine the TE libraries, consensus sequences that showed no full-length fragments (i.e., fragments covering more than 95% of the consensus sequence) in chromosomes were filtered out, and a subset of 1,186 consensus sequences for chr01, 1,174 for chr02 and 1,360 for chr03 were selected. This routine process reduced the complexity of the consensus libraries without any significant loss of global TE coverage. The three resulting libraries were pooled and, after removing redundancy and filtering out those sequences classified as “Potential Host Genes” because they contain host gene Pfam domains, we obtained a final consensus library of 2086 sequences.

Finally, the resulting combined, filtered, and non-redundant library was used to annotate the whole Cupressus genome assembly using the TEannot pipeline.

### Structural and functional annotations

The genetic models were predicted using the Eugene-EP v2.1 pipeline (Carrere et al. 2023), which performs probabilistic sequence model training, genome masking, calculation of transcripts and protein alignments, and integrative gene modeling by EuGene.

Structural annotation of the genes was carried out on the complete genome relying on the previously described set of Isoseq transcripts and tree protein databases: SwissProt, TrEMBL (plants only), and predicted genome proteins of plant models. Moreover, we used the specific TE prediction set described in the previous section for the genome masking step. Default settings were used for this annotation, except for the minimum exon length set to 1bp and maximum intron size increased to 150kb.

To evaluate the accuracy of the 42,980 protein-coding sequences, we assigned a score between 0 and 1 to each protein using Psauron V1.0.4 (Sommer et al. 2025).

Functional annotation was performed on the 42,980 predicted proteins using InterProScan v.5.64-96.0 (Quevillon et al. 2005) for domain and motif searches. We also used the BLASTKoala webservice (http://www.kegg.jp/blastkoala/, September 2024) (Kanehisa et al. 2016) to assign KEGG orthology based on KEGG pathways. The eggNOG-mapper V2.1.12 (Cantalapiedra et al. 2021) was used together with the eggNOG v5 genomes and functional databases of ortholog groups (OGs) (Huerta-Cepas et al. 2019). Finally, the Ensembl Enzyme Prediction Pipeline (E2P2) V.00de829 identified the enzyme codes associated with the 42,980 proteins (https://github.com/carnegie/E2P2) (Chae et al. 2014).

### Annotation of endogenous caulimovirid elements

We searched for endogenous caulimovirid elements (ECVs) in the C. sempervirens genome using the CAULIFINDER package (Vassilieff et al. 2022). The branch A (sequence retriever) of CAULIFINDER was used with default settings to construct a library of consensus sequences that are representative of repetitive ECV sequences present in the genome assembly. This library was then compared to the whole genome using RepeatMasker (Smit, AFA, Hubley, R & Green, P. RepeatMasker Open-4.0. 2013-2015 <http://www.repeatmasker.org>) to annotate the matching ECV copies. The branch B (marker miner) of CAULIFINDER was used to analyse the diversity of Caulimoviridae genera found in the form of ECVs in the C. sempervirens genome assembly and in nine other gymnosperm genome assemblies presented in Fig. S3. The initial step in this process involves the detection of endogenous caulimovirid sequences encoding the reverse-transcriptase (RT) domain and the selection of representative sequences. The selected RT domains are then aligned with a set of reference RT sequences from Caulimoviridae, Retroviridae, and Metaviridae (Ty3/Gypsy LTR retrotransposons), the latter two families representing an outgroup. The alignment is finally used to build a phylogenetic tree.

### LTR retrotransposons analysis

To characterize the dynamics of LTR-retrotransposons in *C. sempervirens* and compare them with *Zea mays*, we first identified intact LTR elements using LTRharvest (El Baidouri and Panaud 2013), filtered for tandem repeats and gag-pol domain presence, and classified sequences into families via SiLiX clustering. Insertion times were estimated by aligning 5′ and 3′ LTRs with MAFFT, calculating Kimura distances using distmat, and applying a substitution rate of 1.3 × 10⁻⁸ substitutions/site/year. Genomic LTR density was approximated by counting reverse transcriptase (RT) domains per Mbp in 1,000-nt sliding windows (via seqkit and BLASTx against a CDD-derived RT database). All commands, scripts (including the custom Perl script for dating), and intermediate datasets are provided in the supplementary repository, alongside genome assemblies and annotations.

We measured the solo-to-intact LTR ratio from the Chr03 of the *Cupressus* genome assembly. We first used LTRharvest in parallel v1.1 from the EDTA package v2.2.1 to collect full length LTR retrotransposons sequences. The resulting library was then processed using the LTR_retriever package v3.0.1. Intact elements were counted using the package’s intact_finder_coarse.pl script, yielding 25,824 copies. Next, solo LTRs were counted using the find_LTR.pl and solo_finder.pl scripts (both part of the LTR_retriever package), resulting in 72,703 solo LTRs. The final ratio was calculated using the solo_intact_ratio.pl script.

### Isoseq analysis

Tools included in the PacBio Isoseq3 distribution were used in the pipeline analysis (https://github.com/ylipacbio/IsoSeq3) with the computer farm facilities of GenoToul (https://doi.org/10.15454/1.5572369328961167E12). For each IsoSeq PacBio library ccs (version 6.2.0 with default parameter) was used to generate one representative consensus sequence from alignment between subreads taken from each ZMWs (zero-mode waveguides). Then, identification of barcodes and removal of cDNA primers and unwanted combinations was performed using lima (version 2.6.0 using --isoseq mode) and chimeric sequences removed using the tool refine wrapped into isoseq3 (version 3.7.0. with default parameter). Five bam files of full-length non-concatemers sequences were created corresponding to each sequencing library. These files were used for de novo transcript reconstruction using cluster wrapped into isoseq3 (version 3.7.0. with default parameter); isoseq3 cluster was used with parameters --verbose and --use-qvs to generate polished High Quality consensus transcript sequences (post-correction accuracy ≥ 99%). The transcripts were aligned to the genome using minimap2 (version 2.24; using -ax splice:hq -t 30 -uf --secondary=no as parameters) and well-mapped sequences were further collapsed to unique transcripts using tama (Kuo et al. 2020) and python script tama_collapse.py with parameters -p prefix -x no_cap.

### Calling and annotation of long non-coding RNA

IsoSeq reads were splice aligned on the genome fasta file indexed with Cse.20240125.gtf gene annotation, produced from Cse.20240125.gff3 with gffread v0.12.6. The alignment was performed with gsnap version 2021-12-17. The alignment file was sorted, compressed and indexed with samtools sort and index version 1.14 with default parameters. Transcripts were called from the alignment file with cufflinks version 2.2.1 using default parameters (https://github.com/cole-trapnell-lab/cufflinks). The resulting GTF transcripts file was filtered to keep only novel genes and transcripts with an in-house awk script retaining only transcripts coming from CUFF genes. The generated GTF file was processed with FEELnc version 0.2 to extract lncRNAs (https://github.com/tderrien/FEELnc). The lncRNAs were annotated with the FEELnc_classifier.pl script.

### Evolutionary history of the conifer genomes from comparative genomics and ancestral genome reconstruction

The ancestral karyotype consists in a ‘median’ or ‘intermediate’ genome made of a clean reference gene order common to the extant investigated species, and derives evolutionary scenarios as described in details Pont et al. 2019, Murat et al. 2017 and Siguret et al. 2026). Briefly, in the first step, conserved gene families (OrthoGroups, OGs) were identified based on the all-against-all BLASTp (Altschul et al. 1990) and the orthology inference tool Orthofinder (Emms and Kelly 2015, 2019). The second step consists of clustering (or chaining) groups of conserved and duplicated genes into ancestral protochromosomes (also referred to as CARs for Contiguous Ancestral Regions) corresponding to independent sets of modern genomic regions from the different investigated species sharing paralogous and/or orthologous relationships, then defining protochromosomes. In the third step, conserved genes (or conserved groups of gene-to-gene adjacencies) between the investigated species within CARs (protochromosomes) are then considered as potentially ancestral genes, defining protogenes. The final step is then performed with dotplots, detecting syntenic regions between modern species and the reconstructed ancestor, and validating that each ancestral protochromosome (or CAR) ‘merges’ or ‘integrates’ all identified conserved (*i.e*. syntenic) regions between modern genomes, with no synteny with any other CARs. From the reconstructed ancestral karyotypes, (i) an evolutionary scenario can then be inferred taking into account the fewest number of genomic rearrangements (including inversions, deletions, fusions, fissions, translocations) which may have operated between inferred ancestor and the modern genomes, and (ii) modern chromosome painting is performed in illustrating ancestral protochromosomes (or CARs) locations in the modern chromosomes based on the syntenic blocks between ancestral and modern genomes, and (iii) nopy number variation of genes was identified on filtered dataset of (551) OGs conserved between the eight gymnosperm-conifer species with MUSCLE (Edgar 2004) and the IQ-TREE 2 (Minh et al. 2020) softwares for clustering inference.

## DATA AVAILABILITY

The C. sempervirens haploid genome assembly and WGS sequencing data have been deposited in the NCBI database under project accession: PRJNA833966.

Full-length transcript sequences obtained from the five PacBio Isoseq3 libraries are available in the SRA bioproject PRJNA836501 (https://www.ncbi.nlm.nih.gov/bioproject/PRJNA836501). The Transcriptome Shotgun Assembly project has been deposited at DDBJ/EMBL/GenBank under the accession GLGR00000000. The version described in this paper is the first version: GLGR01000000.

Genetic maps available in the supplemental data of Hasegawa et al. 2018.

Intermediate analysis for Genomic, Transposable elements and Functional annotation are available in the INRAe public repository (https://entrepot.recherche.data.gouv.fr/dataverse/inrae).

DOIs are provided in the following Table.

Sequences of the contigs mapped in the linkage map and used to validate genome assembly are not publicly available.

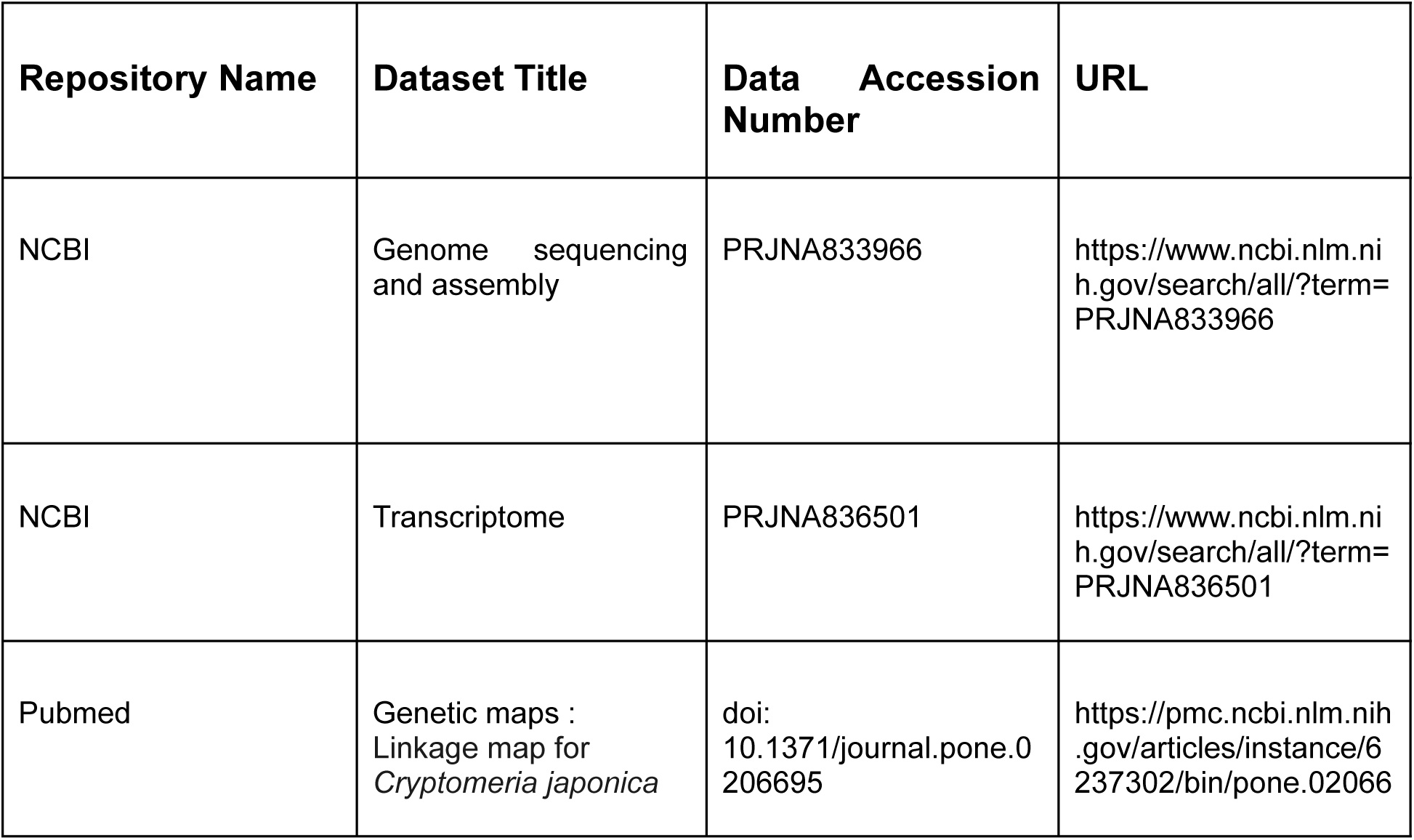
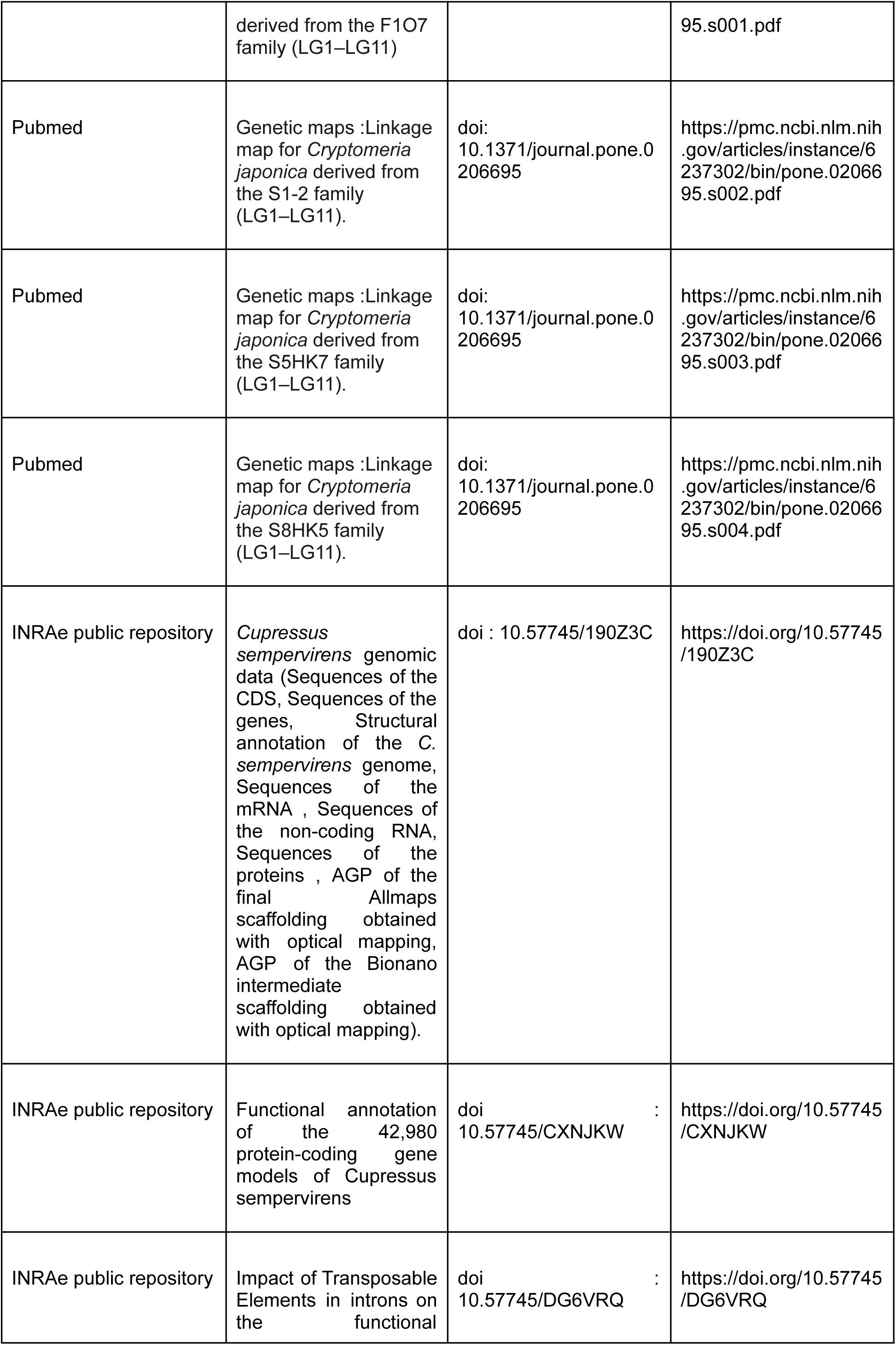
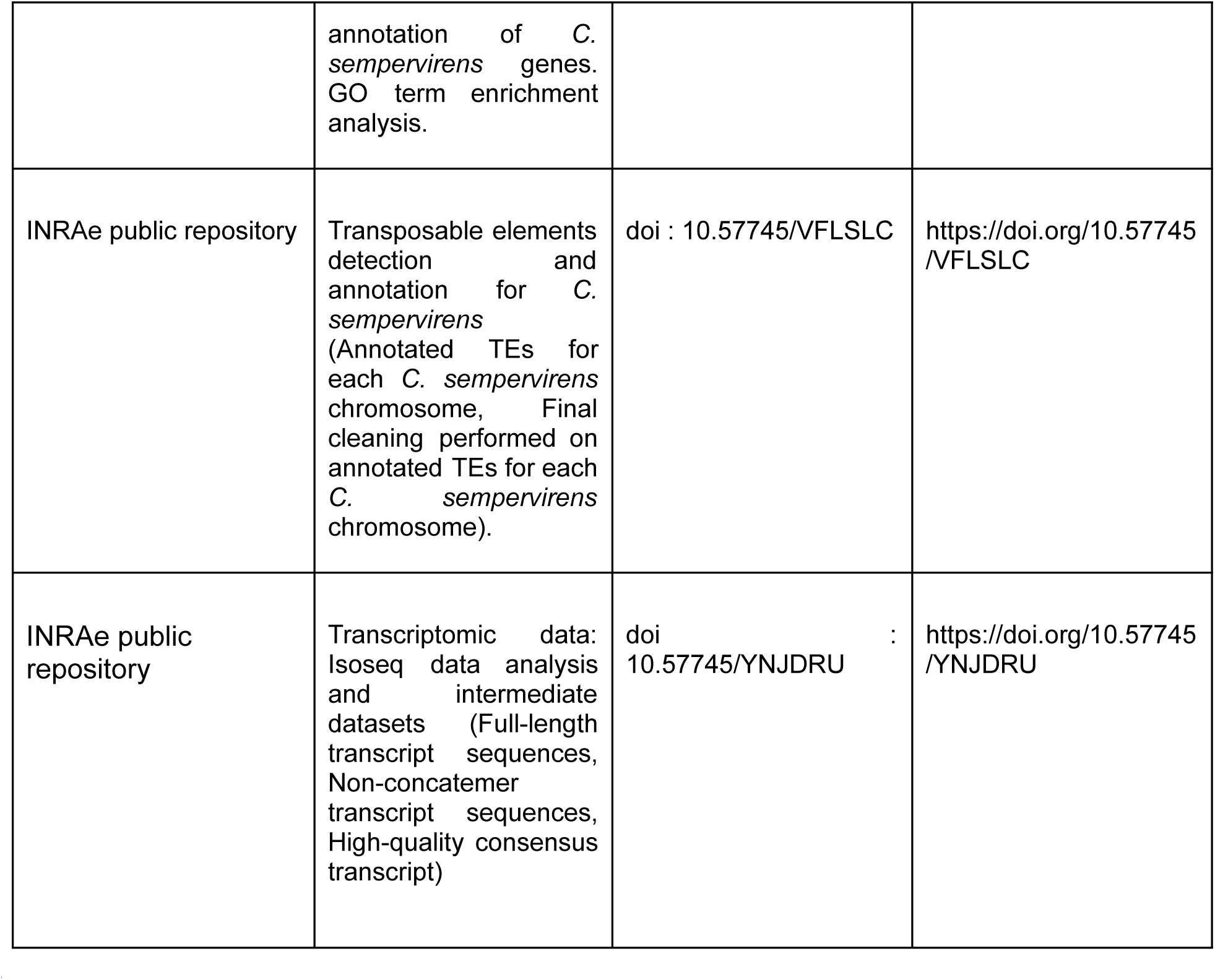

## ABBREVIATIONS

bp: Base pair
BUSCO: Benchmarking Universal Single-Copy Orthologs
CAR: Contiguous Ancestral Region
CCS: Circular Consensus Sequencing
CDS: Coding Sequence
CNV: Copy Number Variation
DAPI: 4′,6-diamidino-2-phenylindole
ECV: Endogenous Caulimovirid Elements
Gb: Gigabase
GO: Gene Ontology
HiFi: High-Fidelity (PacBio)
HMW: High Molecular Weight
kb: Kilobase
lncRNA: Long Non-Coding RNA
LTR: Long Terminal Repeat
LTR-RT: Long Terminal Repeat Retrotransposon
Mb: Megabase
MYA: Million Years Ago
OG: Ortholog Group
PacBio: Pacific Biosciences
PFGE: Pulsed Field Gel Electrophoresis
RT: Reverse Transcriptase
SMRT: Single Molecule Real-Time
TE(s): Transposable Element(s)
uHMW: Ultra-High Molecular Weight
UR: Unequal Recombination
WGD: Whole-Genome Duplication
ZMW: Zero-Mode Waveguide

## AUTHOR’S CONTRIBUTION

AB, CP, JS, CPi, AR, VGi, CPo and WM conceptualized, supervised and coordinated the project. CPi produced the haploïd line of *C. sempervirens*. WM extracted HMW DNA, produced optical maps, hybrid scaffolding and intron studies. JCL extracted RNA and analysed the isoSeq data. CC assembled, scaffolded with genetic maps, annotated structurally the genome and performed telomere analysis. IL produced the functional annotation and regrouped all the data on the dataverse. OP produced the LTR evolution analysis. FM and HV annotated ECV. FM performed the soloLTR analysis. NC realised the TE annotation. VM and FE produced the consensus genetic map. VG and EB realized the HiFi and IsoSeq PacBio library. VP prepared the PacBio data. MDS, CH and JS contributed to the paleogenomic analysis. CK produced lncRNA annotation. All authors discussed the results and contributed to the final manuscript.

## ACKNOWLEDGEMENTS

We are grateful to the genotoul bioinformatics platform Toulouse Occitanie (Bioinfo Genotoul, https://doi.org/10.15454/1.5572369328961167E12) for providing help,computing and storage resources.

We gratefully acknowledge the funding support of INRAE Ecodiv Division. We would like to thank Dr. Saneyoshi Ueno at Forestry and Forest Products Research Institute (Japan) for providing the *Cryptomeria Japonica* consensus linkage map before it was published in 2024 in https://doi.org/10.1186/s12864-024-10929-4.

We thank Jean Thevenet for contribution to plant production and growing, Anne Roig and Touhami Nasradin for their help in tissue sampling for RNAseq analysis.

URGI benefits from the support of Saclay Plant Sciences-SPS (ANR-17-EUR-0007) and the PlantBioinfoPF platform.

During the preparation of this manuscript, the CNRS AI agent, Emmy, has been used for language improvement. All outputs from this tool were critically reviewed, edited, and approved by the authors to ensure accuracy, integrity and originality.

