## Supplemental Data for "Chromosome-scale assembly of the *Cupressus sempervirens* genome unravels new insights into the evolutionary history of conifers"

### SUPPLEMENTARY FIGURES

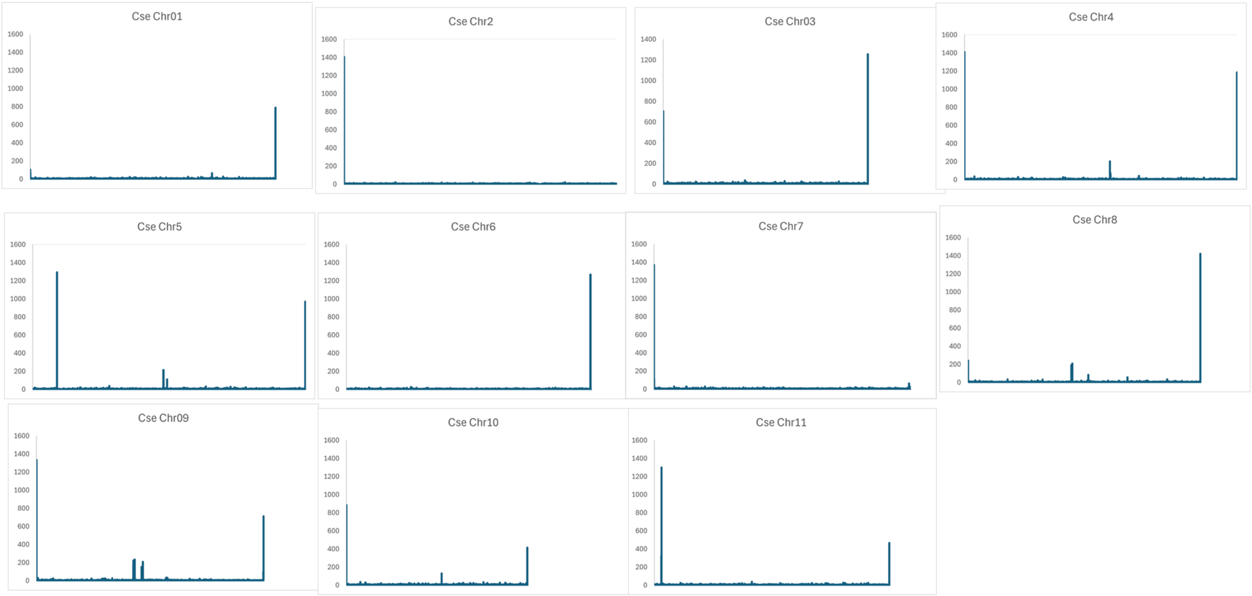


#### Figure S1: Telomere analysis

Chromosome sequences (x axis) are scanned with a telomeric motif and occurrence are presented in y axis

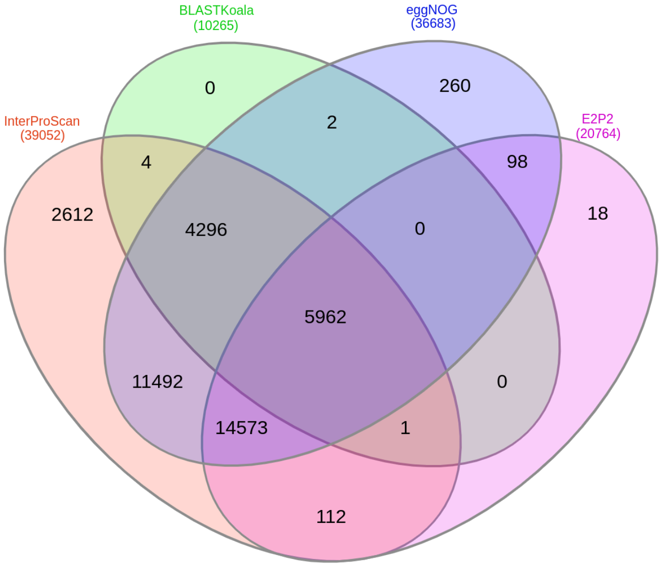


**Figure S2: Functional annotation of the 42,980 C. sempervirens genes.**

Venn diagram of the functional annotation of the *C. sempervirens* genes in Interproscan, KEGG (Kyoto Encyclopedia of Genes and Genomes) with BLASTKoala, eggNOG and E2P2 (Ensembl Enzyme Prediction Pipeline) databases. InterProScan annotation revealed that 39,052 (90.86%) of the *Cupressus* proteins possess functional annotations. Additionally, functional assignments were provided by BLASTKoala and eggNOG for 10,265 (23.88%) and 36,683 (85.35%) proteins, respectively. The E2P2 pipeline further annotated 20,764 proteins (48.31%). In summary, a total of 39,430 (91.74%) of the predicted *Cupressus* proteins received a functional annotation.

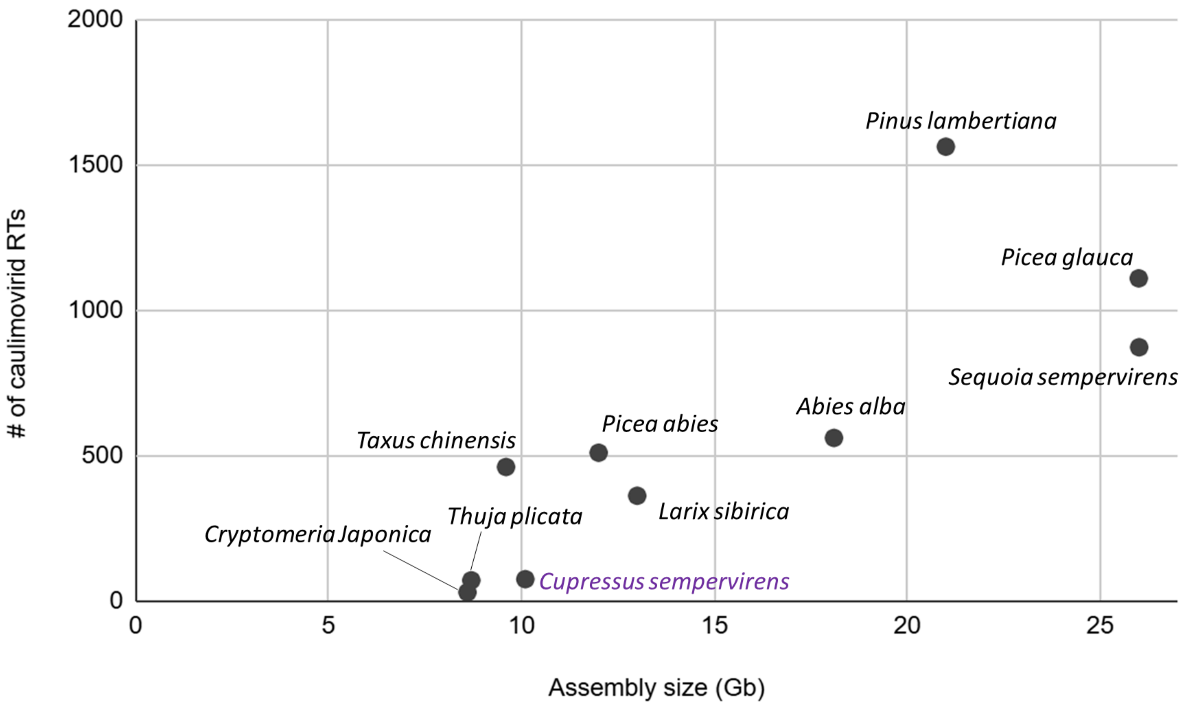


#### Figure S3: Quantification of ECVs in gymnosperm genomes

Scatter chart representing the number of caulimovirid RT domains found in ten gymnosperm genomes as a function of their genome assembly size.

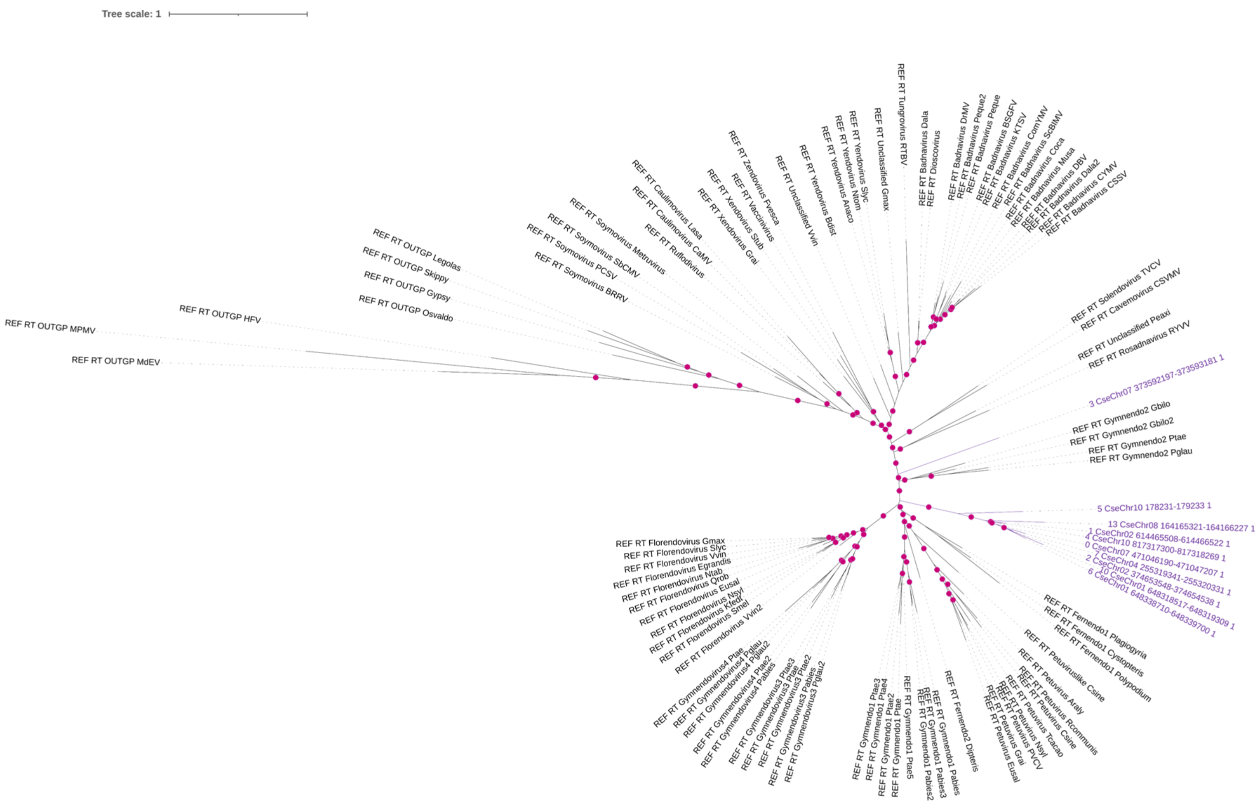


#### Figure S4: Phylogeny of caulimovirids

Phylogenetic tree of the RT domain generated by branch B of CAULIFINDER. The labels of the reference sequences are shown in black; the labels of the reference outgroup sequences contain the tag "OUTGP". The representative *C. sempervirens* caulimovirid RT sequences are shown in purple. The bootstrap values above 70% are indicated as magenta dots.

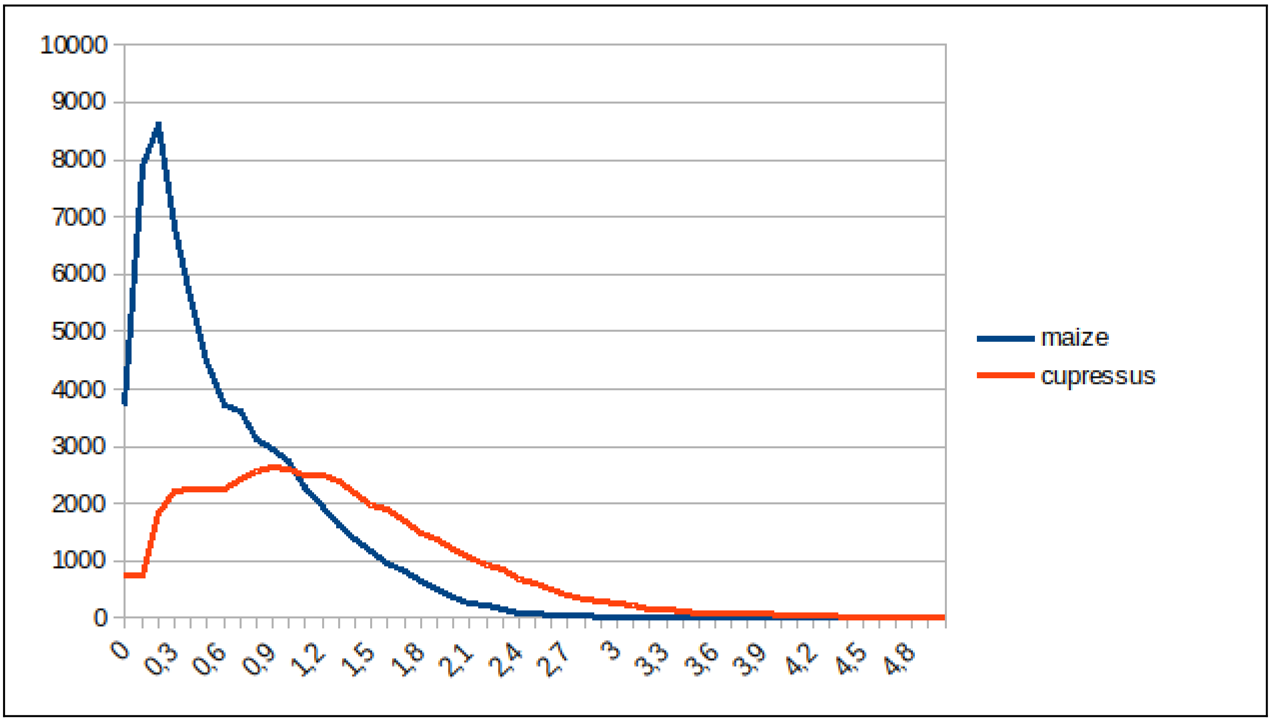


**Figure S5: Age distribution of LTR retrotransposons in the genome of *Cupressus* (red) and maize (blue).**

Age in million years is given on X axis.

### SUPPLEMENTARY TABLES

#### Table S1: Length of the 11 pseudomolecules assembled within Cupressus

| Assembly | Chromosome | Length |
|---|---|---|
| Cupressus.Sempervirens.scaffolded.fasta | CseChr01 | 899 650 123 |
| | CseChr02 | 784 025 100 |
| | CseChr03 | 1 125 945 800 |
| | CseChr04 | 919 939 343 |
| | CseChr05 | 1 059 607 418 |
| | CseChr06 | 717 105 478 |
| | CseChr07 | 938 233 075 |
| | CseChr08 | 852 141 386 |
| | CseChr09 | 832 263 540 |
| | CseChr10 | 995 399 150 |
| | CseChr11 | 861 525 933 |
| Cupressus.Sempervirens.unplaced.fasta | CseChr00c100005_01 | 3007484 |
| | CseChr00c100012_05 | 1531691 |
| | CseChr00s100001_01 | 8745774 |
| | CseChr00s100002_01 | 788974 |
| | CseChr00s100002_03 | 9137269 |
| | CseChr00s100013_2_01 | 2287694 |
| | CseChr00s100103 | 766187 |

#### Table S2: Genome structural annotation metrics with Eugene pipeline

| | | |
|---|---|---|
| Genome | Total number of genes | 77322 |
| | Number of protein coding genes | 42980 |
| | Number of gene with utr both sides | 20728 |
| Genes | % genes with introns | 62 |
| | % genes with five UTR | 51 |
| | % genes with three UTR | 53 |
| | Mean gene length (bp) | 18 300.74 |
| | Total gene length bp | 786 566 072 |
| | Shortest gene bp | 79 |
| | Longest gene bp | 412 140 |
| | Number of single exon gene | 16352 |
| Exons | mean exons per gene | 3.71 |
| | Total exon length bp | 64 229 077 |
| | mean exon length bp | 402 |
| | Longest exon bp | 18759 |
| Introns | Mean intron per gene | 2.71 |
| | mean intron length bp | 6202 |
| | Total intron length bp | 722 336 995 |
| | Shortest intron into gene bp | 40 |
| | Longest intron into gene bp | 132385 |
| CDS | Mean length (bp) | 1062.82 |
| 5'UTR | Mean length (bp) | 325.87 |
| 3'UTR | Mean length (bp) | 503.90 |
| Non protein coding genes | Number of non protein coding genes | 34342 |
| | Mean ncRNA gene length (bp) | 235.64 |
| | Per cent ncRNA genes with introns | 0 |
| Inter protein-coding genes | Mean length | 119320.65 |

#### Table S3: Repeats coverage per chromosome

| Chromosome number | Chromosome size (bp) | Repeats coverage (%) |
|---|---|---|
| Chr01 | 899650123 | 83.53 |
| Chr02 | 784025100 | 82.99 |
| Chr03 | 1125945800 | 81.48 |
| Chr04 | 919939343 | 82.98 |
| Chr05 | 1059607418 | 82.82 |
| Chr06 | 717105478 | 82.48 |
| Chr07 | 938233075 | 83.11 |
| Chr08 | 852141386 | 83.56 |
| Chr09 | 832263540 | 83.11 |
| Chr10 | 995399150 | 82.87 |
| Chr11 | 861525933 | 83.57 |
| ChrUnk | 25498886 | 82.86 |

#### Table S4: Synteny relationships between Gymnosperm-conifer complex genomes.

The table provides, for the investigated species (in columns) and ancestral chromosomes ('AncChr', in lines), the associated chromosomal synteny relationships with the chromosomes involved and the number of orthologous genes in parenthesis

| | **Sequoiadendron_giganteum** | **Cupressus_sempervirens** | **Taxus_chinensis** |
|---|---|---|---|
| 1 | chr6 (429), chr9 (281) | chr4 (362), chr8 (265) | chr2 (236), chr4 (313) |
| 2 | chr4 (433), chr7 (292) | chr1 (256), chr2 (376) | chr3 (318), chr8 (219) |
| 3 | chr2 (21), chr11 (499) | chr2 (248), chr9 (204), chr10 (25) | chr5 (227), chr9 (244) |
| 4 | chr1 (136), chr3 (383) | chr3 (489) | chr1 (70), chr7 (322) |
| 5 | chr3 (385) | chr3 (378) | chr1 (295), chr6 (33), chr10 (30) |
| 6 | chr10 (386) | chr6 (330) | chr4 (327) |
| 7 | chr7 (353), chr9 (27) | chr1 (321), chr8 (26) | chr2 (315) |
| 8 | chr8 (340) | chr11 (336) | chr5 (99), chr6 (174) |
| 9 | chr10 (348) | chr6 (325) | chr5 (295) |
| 10 | chr1 (334) | chr7 (288) | chr11 (253) |
| 11 | chr2 (322) | chr10 (293) | chr12 (228) |
| 12 | chr4 (258) | chr9 (258) | chr1 (259) |
| 13 | chr9 (265) | chr8 (248) | chr7 (236) |
| 14 | chr8 (268) | chr11 (252) | chr7 (32), chr10 (188) |
| 15 | chr5 (265) | chr5 (263) | chr3 (226) |
| 16 | chr4 (28), chr5 (52), chr12 (211) | chr5 (214), chr9 (18) | chr11 (213) |
| 17 | chr4 (43), chr5 (169) | chr5 (163), chr9 (37) | chr8 (167) |
| 18 | chr6 (196) | chr4 (169) | chr6 (151) |
| 19 | chr1 (185) | chr7 (177) | chr10 (153) |
| 20 | chr2 (152) | chr10 (136) | chr9 (135) |
| 21 | chr2 (131) | chr10 (118) | chr5 (19), chr6 (84) |

| | **Pinus_tabuliformis** | **Ginkgo_biloba** | **Cycas_panzhihuaensis** |
|---|---|---|---|
| 1 | chr9 (722) | chr10 (534) | chr2 (219), chr7 (345) |
| 2 | chr4 (798) | chr8 (572) | chr1 (366), chr9 (188) |
| 3 | chr7 (349), chr10 (272) | chr1 (80), chr2 (262), chr12 (116) | chr2 (170), chr7 (122), chr9 (208) |
| 4 | chr10 (308), chr12 (242) | chr4 (255), chr11 (142) | chr3 (166), chr4 (148), chr6 (69), chr10 (83) |
| 5 | chr1 (127), chr3 (337) | chr7 (308) | chr5 (328) |
| 6 | chr6 (432) | chr1 (319) | chr10 (313) |
| 7 | chr3 (423) | chr6 (310) | chr2 (288) |
| 8 | chr12 (387) | chr4 (273) | chr4 (303) |
| 9 | chr11 (398) | chr3 (303) | chr11 (271) |
| 10 | chr5 (354) | chr1 (269) | chr8 (276) |
| 11 | chr5 (314) | chr12 (232) | chr9 (245) |
| 12 | chr2 (344) | chr5 (225) | chr1 (102), chr4 (19), chr6 (43), chr11 (72) |
| 13 | chr8 (360) | chr9 (224) | chr6 (241) |
| 14 | chr11 (318) | chr9 (228) | chr1 (93), chr10 (112) |
| 15 | chr8 (311) | chr2 (211) | chr1 (141), chr6 (71) |
| 16 | chr1 (334) | chr6 (190) | chr2 (196) |
| 17 | chr2 (292) | chr5 (173) | chr1 (54), chr3 (57), chr8 (69) |
| 18 | chr7 (238) | chr3 (158) | chr7 (78), chr11 (73) |
| 19 | chr1 (228) | chr7 (139) | chr5 (151) |
| 20 | chr6 (181) | chr11 (114) | chr3 (134) |
| 21 | chr6 (122) | chr11 (91) | chr3 (99) |

#### Table S5: Genome sources used for paleogenomic analysis

| Species name | Source |
|---|---|
| *Pinus lambertiana* | https://treegenesdb.org/FTP/Genomes/Pila/v1.5/ |
| *Picea abies* | https://treegenesdb.org/FTP/Genomes/Paab/ |

| | |
|---|---|
| *Picea glauca* | http://congenie.org/ |
| *Abies alba* | https://treegenesdb.org/FTP/Genomes/Abal/v1.1/ |
| *Larix Sibirica* | https://ftp.ncbi.nlm.nih.gov/genomes/all/GCA/004/151/065/GCA_004151065.1_LarixSibirica0.1/GCA_004151065.1_LarixSibirica0.1_genomic.fna.gz |
| *Sequoia sempervirens* | https://treegenesdb.org/FTP/Genomes/Sese/v2.1/ |
| *Thuja plicata* | https://phytozome-next.jgi.doe.gov/info/Tplicata_v3_1 |
| *Cupressus sempervirens* | This study |
| *Cryptomeria japonica* | https://ftp.ncbi.nlm.nih.gov/genomes/all/GCA/027/924/625/GCA_027924625.1_CJA_r1.0/GCA_027924625.1_CJA_r1.0_genomic.fna.gz |
| *Taxus chinensis* | https://db.cngb.org/search/assembly/CNA0036166/ |
